# GenomeGraphR: A user-friendly open-source web application for foodborne pathogen Whole Genome Sequencing data integration, analysis, and visualization

**DOI:** 10.1101/495309

**Authors:** Moez Sanaa, Régis Pouillot, Francisco Garces Vega, Errol Strain, Jane M. Van Doren

**Affiliations:** Center for Food Safety and Applied Nutrition, Food and Drug Administration, College Park, Maryland, United States of America

**Author notes:** Current Address: Food Safety Risk Assessment Unit, Risk Assessment Department, ANSES (Food agency for food, environmental and occupational health & safety), Maisons-Alfort, France (Moez Sanaa).

**Keywords:** WGS, foodborne pathogen, network analysis, SNP distance, software

## Abstract

Food safety risk assessments and large-scale epidemiological investigations have the potential to provide better and new types of information when whole genome sequence (WGS) data are effectively integrated. Today, the NCBI Pathogen Detection database WGS collections have grown significantly through improvements in technology, coordination, and collaboration, such as the GenomeTrakr and PulseNet networks. However, high-quality genomic data is not often coupled with high-quality epidemiological or food chain metadata. We have created a set of tools for cleaning, curation, integration, analysis and visualization of microbial genome sequencing data. It has been tested using *Salmonella enterica* and *Listeria monocytogenes* data sets provided by NCBI Pathogen Detection (160,000 sequenced isolates). GenomeGraphR presents foodborne pathogen WGS data and associated curated metadata in a user-friendly interface that allows a user to query a variety of research questions such as, transmission sources and dynamics, global reach, and persistence of genotypes associated with contamination in the food supply and foodborne illness across time or space. The application is freely available (https://fda-riskmodels.foodrisk.org/genomegraphr/).

## Introduction

The implementation of Whole Genome Sequencing-based techniques (WGS) as a routine typing method for specific foodborne pathogens has significantly improved surveillance, increased the number of outbreaks being detected, shortened the time to detect them and the time to find their source [1, 2, 3, 4, 5, 6].

Food safety risk assessments and large-scale epidemiological investigations have the potential to provide better and new types of information when WGS data are effectively integrated [7, 8, 9, 10]. For example, WGS data have the potential to characterize the relative importance of sources (attribution) and to identify strains that circulate widely and those that do not. WGS data also have the potential to provide information about how pathogen strains and sources are linked (or not) and how these sources and links persist and evolve over time and space. WGS data can also provide information about the distribution of virulence among strains and which among these strains are most often associated with foodborne illness. Integration of these data into risk assessments and population level epidemiology studies was initially limited by the size, scope, and consistency of available WGS databases because these studies require representative or population level data.

Today, WGS collections such as the NCBI Pathogen Detection (https://www.ncbi.nlm.nih.gov/pathogens) have grown significantly through improvements in technology, coordination, and collaboration of the GenomeTrakr and PulseNet networks in the US. Presently, the two networks sequence and release about 5,000 new bacterial genomes per month [11, 12] which makes their analysis and interpretation increasingly demanding.

Our original goal was to explore what we, as risk assessors and epidemiologists, could learn from the NCBI WGS data. We found, however, that despite extensive efforts by the GenomeTrakr network to collect and standardize isolate metadata, the first thing we had to do was to curate and further standardize the metadata associated with isolates in order to answer the kinds of questions we were interested in. In doing this we developed a new web-based visualization and analytic application, GenomeGraphR. It consists of a Shiny-R web application that allows user-friendly interactive visualization of genomic data and comprises a set of dynamic illustrations including networks, trees, epidemic curves, maps, figures and tables linked to and embedded with cleaned and structured metadata. The application is hosted online as an R Shiny web server application (https://fdariskmodels.foodrisk.org/genomegraphr/). This manuscript describes our research and results to date.

## Materials and methods

### Data source

In 2013, the US Food and Drug Administration (FDA) created a United States-based open-source whole-genome sequencing network of state, federal, international, and commercial partners, GenomeTrakr, that collects and shares genomic and associated metadata from foodborne pathogens [1]. The high-quality genomic data are deposited in public databases at the National Center for Biotechnology Information (NCBI) of the National Institutes of Health (NIH). The NCBI uses an automated pipeline (ftp.ncbi.nlm.nih.gov/pathogen/Methods.txt) which offers quality control procedures and removal of low-quality raw reads, groups the genomes into clusters based on *wg-MLST*, calculates the pairwise single nucleotide polymorphism (SNP) distance for all pairs within the same *cluster* and finally converts the SNP information for each target partition into a maximum compatibility tree. As of August 1^st^, 2018, the network includes sequence data for almost 140,000 *Salmonella enterica*, 51,000 *E. coli* and *Shigella spp*., 20,000 *Listeria monocytogenes* and 20,000 *Campylobacter jejuni* isolates (https://www.ncbi.nlm.nih.gov/pathogens/). In addition, the NCBI provides web-based tools for exploring these public data, including search tools and SNP-based trees (www.ncbi.nlm.nih.gov/pathogens/isolates).

For each pathogen, the clusters are regularly updated by NCBI as more strains are submitted by users. The GenomeGraphR application introduced here is currently populated with *L. monocytogenes* and *S. enterica* data downloaded from ftp://ftp.ncbi.nlm.nih.gov/pathogen/Results/ on July 31^st^, 2018. The *S. enterica* metadata file (distribution #1171) included data from 139,754 strains. *Salmonella* SNP file included 11,576,777 pairwise SNP distance. Similarly, the *L. monocytogenes* database from the same date (distribution #952) included metadata from 18,696 strains and 947,756 pairwise distances.

### WGS metadata curation and organization

A standard minimum set of metadata fields are collected along with each food or environmental isolate submitted by the GenomeTrakr network. Clinical isolates submitted by PulseNet have a delayed release of these standard fields. These standard fields are included in NCBI’s pathogen metadata template for BioSample submission and include the following general contextual (who, what, when, and where) information: who collected the isolate, its taxonomic name, its date of collection (day, month, year), its country/state of origin, and its isolation source (*e.g*., clinical, cilantro, avocado, environmental swab,…). However, two of these metadata fields are entered as free text (“collected by” and “isolation source”). The resulting heterogeneity makes it significantly difficult to query the database efficiently at a large scale or to perform basic statistical analyses on the whole database, as needed in epidemiology investigations or quantitative microbial risk assessment. A simple question such as: “how many *Salmonella* strains have been isolated from produce?” cannot be answered. The first objective of the present work was then to develop and apply a systematic metadata preparation procedure to improve the ability of researchers to exploit the rich data in the current NCBI database.

There are two mandatory fields describing the isolate source. One determines which pathogen attribute package is used and has only two allowed entries, “clinical/host-associated” or “environment/food/other”, and the other is a free text field called “isolation_source”. The heterogeneity of this second field is extreme. For example, if the isolation source was a fish, it could be recorded as “fish”, “smoked trout”, “fresh albacore”,… or by its Latin name, including typos. An R code using perl-like regular expression was developed to curate the *S. enterica* database (5,620 distinct isolation sources in the database) while the smaller *L. monocytogenes* database was handled using a spreadsheet (2,357 distinct isolation sources). To classify isolation sources, we adapted the Interagency Food Safety Analytics Collaboration (IFSAC) scheme [13] which has the advantages of being simple enough that a basic knowledge of the isolation source is in general enough to classify a sample. Moreover, this classification was developed by a consensus of three major US agencies (Centers for Disease Control and Prevention (CDC), FDA and Food Safety and Inspection Service (FSIS)) and is currently used to estimate source attribution from outbreak related foodborne illnesses. The IFSAC scheme is a five-level hierarchy for categorizing foods with increasingly specificity, resulting in a total of 78 food categories. Without altering its main structure, the original IFSAC scheme was adapted and extended to include non-food isolates as well as composite foods. Not all the isolates were sufficiently described to enable a plausible assignment to a specific level. A field giving the “best-known level” was then created.

In addition to the isolation source, we also standardized other important metadata fields such as the date, country, location of sample from which the strain was isolated, host name, project center, etc. Significantly, the date of sample collection was for one category of isolates (i.e., clinical isolates) seldom provided in the current dataset. Because the date of target (WGS) creation in the database was always available (automatically generated within the bioinformatic pipeline), GenomeGraphR gives users the option to select as a proxy for the date of collection (as entered by the strain submitter, with missing data), the date of target creation (no missing data), or the date of collection replaced, if missing, by the date of target collection. However, using the date of target creation could be misleading when analyzing relatedness between strains if strains were sent for WGS analysis long after the date of sample collection (e.g., samples collected during the early 1990’s but not processed with WGS until 2015).

The sampling location information was also frequently missing in the NCBI database. GenomeGraphR gives users the option to select the geographical location where the sample was collected (as indicated by the submitter, with frequent missing data), the location of the submitter (the submitting center is automatically generated by the pipeline and all the submitting centers were geo-localized), or the location of sample collection, replaced, if missing, by the location of the center. When analyzing geographical the WGS data, selecting the center location could be misleading because a laboratory may process isolates from samples collected from all over the world (*e.g*., FDA-Center for Food Safety and Applied Nutrition in Maryland U.S. processing samples from Brazil).

### SNP clusters, SNP threshold

The number of SNPs separating isolate genomes are used to assess the genetic relatedness among isolates [13]. If a small SNP distance is observed, the isolates can be considered as closely related and the likelihood that they arise from a common source is high. When the number of SNPs between two isolates is large, these isolates are considered distantly related, implying that they are probably not originating from the same source or reservoir of the bacteria population. By construction, the NCBI database provides the SNP distance between two isolates only within “SNP clusters”. A SNP cluster is a cluster of isolates where each isolate is less than or equal to 50 SNPs distant from other members of the cluster.

Presently, there is general agreement on what is considered closely related, however the community agrees that using an exact SNP threshold to determine if isolates may be treated as arising from the same source is not advisable [14]. In GenomeGraphR, the user can vary the threshold value and explicitly explore its impact on the result. The SNP threshold is set to 12 by default.

### Network Analysis

The theories, metrics and tools used in network analyses have their foundations in Social Network Analysis (SNA), which uses graph theory principles, computing techniques and resources, to analyze network structures and their interactions [13]. Network Analysis methods are currently used in a variety of domains. In microbiology, they provide tools for tackling a multitude of complex phenomena, including the evolution of gene transfer, composite genes and genomes, evolutionary transitions, and holobionts [14].

We applied Network Analysis to the WGS NCBI data set. The data are organized as a graph. A graph is a set of entities *V*, called vertices or nodes that are connected. The connections between the vertices are called edges or links (*E*). For a given graph *G* = (*V, E*), we denote by *n* = |*V*| its number of nodes, by *m* = |*E*| its number of links, and by *N*(*v*) = {*u* ∈ *V*, (*v, u*) ∈ *E*} the set of neighbors, or neighborhood, of node *v*. In GenomeGraphR, the nodes correspond to isolates and the nodes are connected when their SNP difference is lower than a user specified threshold. The graphs are undirected, *i.e*. no distinction is made between (*u, v*) and (*v, u)*, and weighted using a value equal to the SNP difference between two linked isolates. Nodes are grouped in connected components (CC), that are a subgraph in which any two vertices are connected to each other by paths, and which is connected to no additional vertices in the graph (G). Some topological properties commonly used in social network analysis are calculated to describe the pattern of inter-relationships between the isolates.

### Search and analysis Strategy

GenomeGraphR can be used to address a variety of questions including the following:

- How many (and what fraction of) food isolates from a given food category are related to clinical cases?
- How often are isolates from a specific food category linked to clinical isolates? Is this frequency periodic or sustained?
- What is the global reach of isolates from a given food? How does this compare to the global reach of connected clinical cases?
- How often are strains isolated from food originating from a given geographic area linked to strains isolated from clinical cases occurring the U.S. or other countries?

Queries and data manipulation were pre-designed using igraph [15] and data.table [16] R packages functionalities to collect the relevant information to answer those questions. GenomeGraphR tool offers three ways to select strains to explore: (1) from the isolation source category tree; (2) from the NCBI table including the metadata where the user can filter on one or more fields of the table; and (3) from an uploaded user file containing a predefined set of strains of interest (defined, *e.g*. by their target accession number) that will be matched with the NCBI file. Once the set of strains of interest is defined, the tool provides instantly the numbers of clinical isolates with SNP difference less or equal to the chosen threshold from the selected strains, and the number of corresponding CC.

A “***globe graph***” visualizes the query result. This visualization uses the VisNetwok [17] [18] R package that provides an R interface to the ‘vis.js’ JavaScript charting library and allows an interactive visualization of networks. This first visualization (**[Fig 1**, step B), where only edges between clinical and the selected source are drawn, allows a general overview of the links between strains from the selected source and clinical strains. Each CC could be interpreted as a potential outbreak to be confirmed with further analyses.

**Fig 1:**
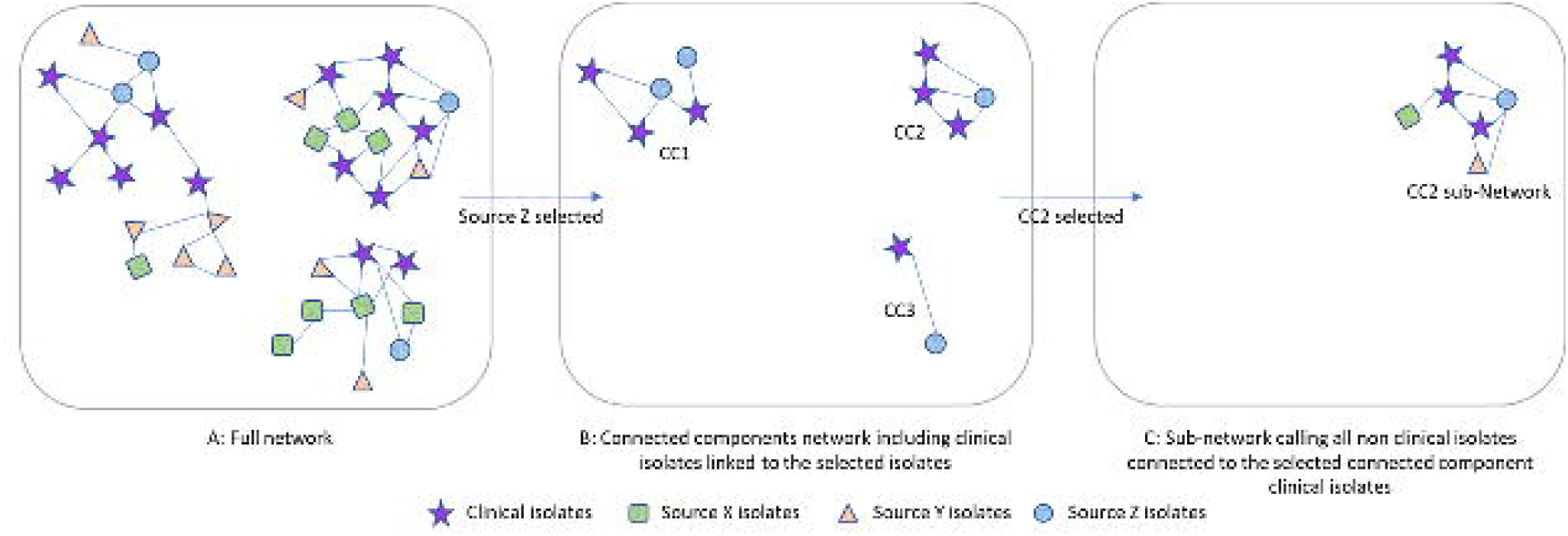
Simplified example of the search strategy. A. the complete network includes all nodes and link the nodes that are closer, in SNP distance, than a given, user-specified, threshold. B. Selecting some strains (e.g. based on their isolation source), the connected components are limited to those strains and the clinical strains closer than the SNP threshold. This graph shows the strains from the isolation source that are potentially linked to clinical strains. It includes only clinical strains and strains from the selected isolation source. C. in order to verify if these links are meaningful, all the additional strains, from any sources, that are closer than the SNP threshold to the clinical strains are recalled, forming a sub-network.

To start further analyses, the user selects one CC of interest from the globe-graph. Once the CC is selected, the tool recalls all non-clinical strains linked to the clinical strains belonging to the selected CC and adds them to the CC to construct a sub-network (**[Fig 1**, step C). In this subnetwork all the links with SNP distance less than or equal to the threshold are drawn. The purpose of this additional analysis is to find and explore other possible linked strains to the clinical strains in the CC. It is performed using dynamic graphical and tree illustrations, embedded with metadata, of the CC strains including minimum spanning tree, classical clustering trees, circular plots, Sankey plots, maps describing the geographical distribution of isolates and epi-curves showing strain occurrence over time.

A ***minimum spanning tree (MST)*** is a subgraph that connects all the nodes (isolates) together, without any cycle and with the minimum possible total link weight (SNP). This path could be interpreted as the most likely chain of pathogen transmission. MSTs are primarily convenient for investigating relationships of microorganisms where not enough genetic diversity has accrued to permit the use of more mathematically sophisticated algorithms for inferring population structure. Prim’s algorithm, implemented in igraph R package, was used to derive the MSTs using the full pairwise SNP difference matrix of the selected sub-network.

#### Clustering trees

Many clustering methods exist in the literature [21]. In our application we are using hierarchical clustering methods implemented in the “hclust” function from R. The input data is a distance matrix which corresponds to the pairwise SNP difference matrix for each sub-network. By default, the complete linkage method is used. However, the user has the option to choose other methods (e.g. Ward’s, centroid, average,…). The dendextend [17] R package was used for extending dendrogram objects provided by “hclust”. Clustering tree analysis is the most common method used for SNP WGS data.

***Circular and Sankey Plots*** were used to visualize the distribution of links between all the categories of isolates, including clinical isolates. For each sub-network, the number of links between and within sources were counted and illustrated graphically.

***Maps*** were created to display available geographical data. The Leaflet R package [19] was used to allow extensive map annotation and dynamics over time. However, this descriptive analysis was limited by the absence of geographical localization (U.S. states) for most of the clinical cases and for some of the non-clinical isolates. To prevent misinterpretation when geographical data are missing (primarily the case for clinical isolates where only country level is available) the isolates are displayed as markers in a predefined polygon.

An ***epi-curve*** occurrence of isolates over time, is provided using different possibilities for the time scale. The timeline shows strain isolation/submission events (as selected) for all isolates in the CC.

#### Data download

All the data queried and used in the previously described illustrations can be downloaded from the download menu. Moreover, each illustration (map, network, sub-network, MST, epi-curve, or trees) can be saved in PNG format.

## Results

### Metadata Cleanup and Organization

While a minimum set of metadata field requirements are a progressive step, in instance the isolation sources are currently entered as non-controlled free text, which required time-consuming verification and validation procedures before being integrated with genomic data for analyses. Moreover, public health agencies have different constraints about the level of metadata that can be made openly accessible. For example, the Centers for Disease Control and Prevention (CDC) provide only the years of clinical cases occurrence and does not communicate the geographical location of the cases. GenomeGraphR integrates NCBI metadata that has been cleaned and organized. We used a hierarchical classification/categorization of isolation sources built on the IFSAC scheme [21], chosen for its simplicity, acceptability, and use in the food safety attribution domain.

A total of 139,754 isolates of *S. enterica*. were submitted to NCBI from 2010 to 2018 as of July 31^st^, 2018. The isolation source of 812 (0.6 % of all the strains) were not classified because of missing or unclear/unintelligible data. For *L. monocytogenes* only 59 isolates out of a total of 16,567 were not assignable to any of the defined isolate categories. The distribution of isolates by major isolate categories is presented in Table 1. The categorization scheme applied to *L. monocytogenes* and *S. enterica* strains consists of the eight-level hierarchy for categorization of foods developed by IFSAC [21], extended to include environmental and animal (non-food) sources and applied here to strain isolation sources NCBI. [**Fig 2** illustrates the hierarchy for the non-clinical strains and the volume of strains associated with each level using a Sankey plot.

**Table 1:**
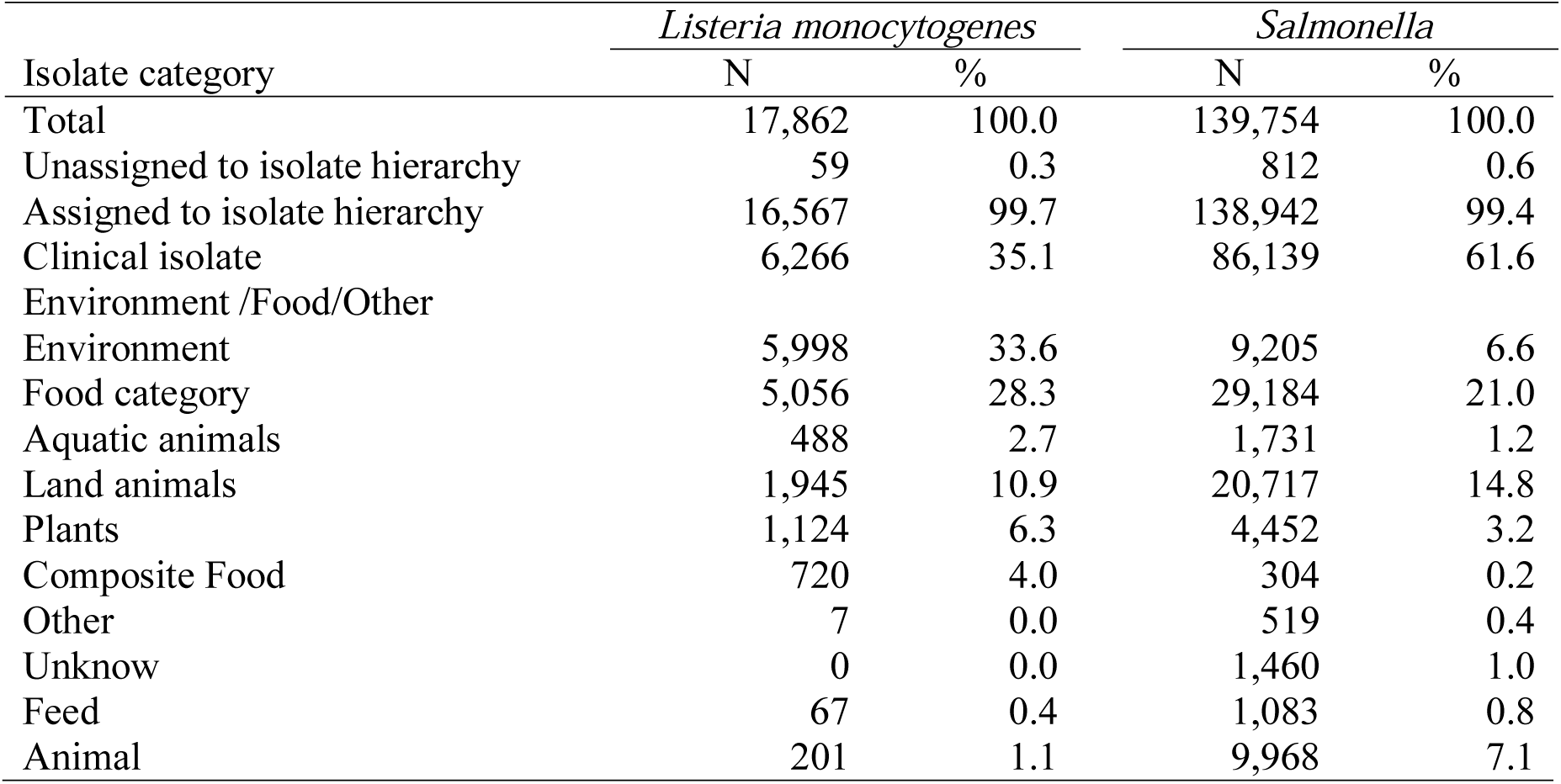
Isolates distribution by major isolate categories

| Isolate category | *Listeria monocytogenes* |  | *Salmonella* |  |
| --- | --- | --- | --- | --- |
|  | N | % | N | % |
| Total | 17,862 | 100.0 | 139,754 | 100.0 |
| Unassigned to isolate hierarchy | 59 | 0.3 | 812 | 0.6 |
| Assigned to isolate hierarchy | 16,567 | 99.7 | 138,942 | 99.4 |
| Clinical isolate | 6,266 | 35.1 | 86,139 | 61.6 |
| Environment /Food/Other |  |  |  |  |
| Environment | 5,998 | 33.6 | 9,205 | 6.6 |
| Food category | 5,056 | 28.3 | 29,184 | 21.0 |
| Aquatic animals | 488 | 2.7 | 1,731 | 1.2 |
| Land animals | 1,945 | 10.9 | 20,717 | 14.8 |
| Plants | 1,124 | 6.3 | 4,452 | 3.2 |
| Composite Food | 720 | 4.0 | 304 | 0.2 |
| Other | 7 | 0.0 | 519 | 0.4 |
| Unknow | 0 | 0.0 | 1,460 | 1.0 |
| Feed | 67 | 0.4 | 1,083 | 0.8 |
| Animal | 201 | 1.1 | 9,968 | 7.1 |

**Fig 2:**
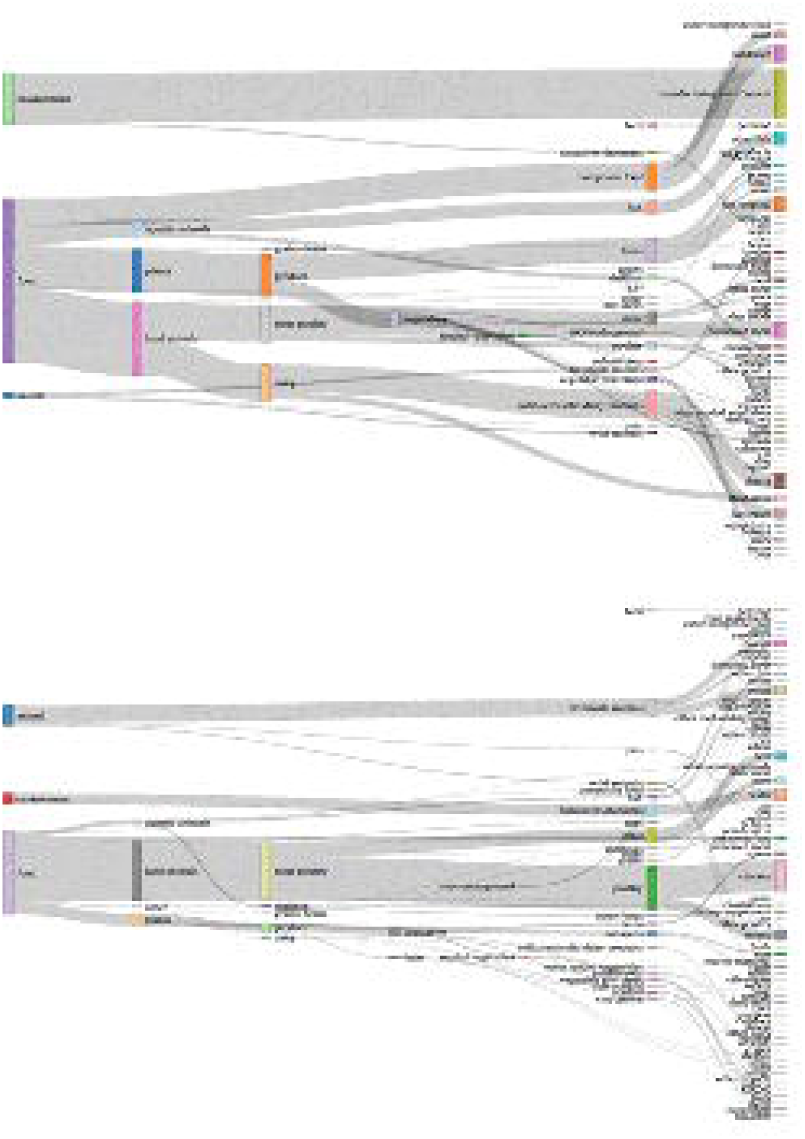
Categorization scheme of strain isolation sources with relative numbers of isolates illustrated by the width of the bank in this Sankey plot– Non-clinical strains. Top: *L. monocytogenes* strains (root: 10,912 isolates), Bottom: *S*. *enterica* strains (root: 49,525 strains).

The process of controlled vocabulary cleanup conducted in this study is time consuming and costly, despite an effort made to manage data cleanup through automatization and coding procedures. However, without this effort more than a quarter of records returned in a keyword search would be lost. As an example, filtering the keyword “fish” before metadata cleanup returned 396 records for *S. enterica*, while, after the use of our controlled vocabulary, the same query returned 949 records.

### Network characteristics

Using the NCBI curated data in GenomeGraphR, we characterize the pathogen networks to provide insights into the extent to which each isolate collection represents the full range of circulating pathogen strains. The number of isolates submitted to the GenomeTrakr network each year continues to increase for both the *L. monocytogenes* and *S. enterica* collections ([**Fig 3**). The SNP-clustering outputs performed by NCBI, grouped the 18,696 *L. monocytogenes* strains into 1,752 SNP-clusters and the 139,754 *S. enterica* strains in 6,282 SNP-clusters. However, 3,844 and 14,415 strains of *L. monocytogenes* and *S. enterica* were not assigned to a SNP cluster because the SNP-analysis found no strain within a SNP difference less or equal to 50 for them. For both pathogens, the total number of SNP-clusters continues to increase over time ([**Fig 3**) which means that an important part of the genetic diversity of *S. enterica* and *L. monocytogenes* strains has yet to be captured in the collection.

**Fig 3:**
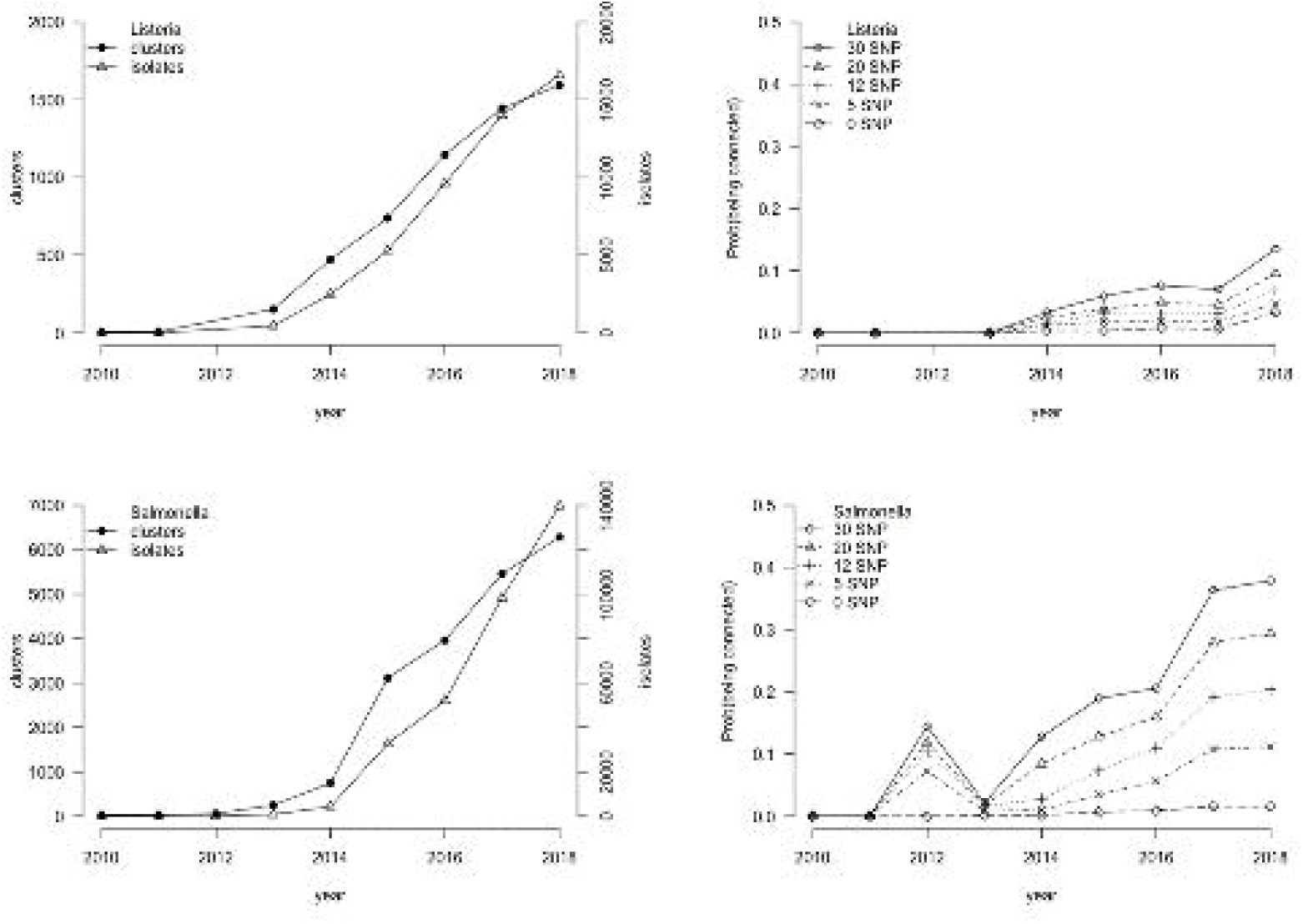
Left: Numbers of isolates (scale on the right axis) and number of SNP clusters (scale on the left axis) as a function of time (creation date of the target in the NCBI database), for *L. monocytogenes* (top) and *S. enterica* (bottom). Right: Probability for a newly created clinical strains of being genetically matched with a non-clinical strain previously isolated, as a function of time and SNP threshold for *L. monocytogenes* (top) and *S. enterica* (bottom). Note: the 2013 artifact for *S. enterica* is linked to the massive inclusion of new strains in 2013.

The evolution over time of the probability of pairing genetically an isolated clinical strain with at least one non-clinical pre-existing strain in the database is presented for different SNP thresholds ([**Fig 3**). This metric characterizes the likelihood that a possible food or environmental source of a foodborne illness associated with that pathogen could be identified immediately from the NCBI database, without collecting additional isolates. Since 2013, this probability has been increasing for both *S. enterica* and *L. monocytogenes* strains because of the increasing size and scope of the collection but remains relatively low (∼15% for *Salmonella* and ∼10% for *L. monocytogenes* in 2018).

The number of connections per strain, called degree, is a characterization of the genetic relatedness of strains in the collection. The change in degree through time provides a measure of the persistence of genetically related strains. The degree was calculated for each strain within the collection with a SNP threshold equal to 12 and for the accumulated strain collections over time ([**Fig 4**). The degree distribution is highly asymmetric, most of the nodes have low degrees while a small but significant fraction of nodes have an extremely high degree. At a SNP threshold of 12, the average degree for *L. monocytogenes* isolates increases between 2013 and 2015 but was approximately constant between 2015 and 2018 (ranging between 21.3 and 24.7). For *S. enterica* isolates, at a SNP threshold of 12, the average degree is increasing over the entire period and peaks at 117 in 2018. This could be easily extended to explore how the degree distribution and mean value vary with SNP distance.

**Fig 4:**
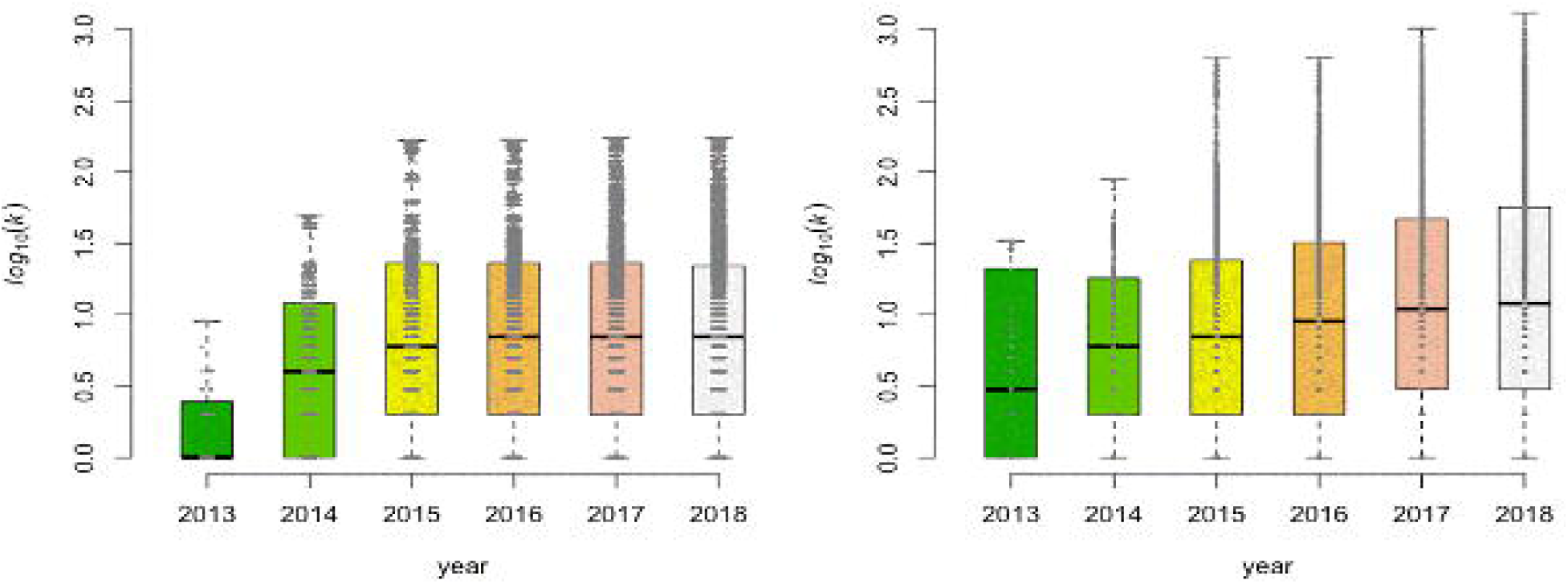
Box-plot of the number of connections per strains (Degree - k) as a function of the year of creation of the target, per year. (left: *L. monocytogenes*; right: *S. enterica*. SNP threshold = 12).

The minimum number of links for a CC with *n* nodes is (n – 1), while the maximum possible number of links is n X (n – 1)/2. A CC with a maximum number of links is called a clique and corresponds to the situation of complete connection between strains belonging to the same CC. The number of connections per CC provides a descriptive metric of its structure. [**Fig 5** shows for both pathogens the high connectivity between strains belonging to the same CC using a SNP threshold of 12: for CC with *n* > 10, the number of links is closer to the maximum than the minimum.

**Fig 5:**
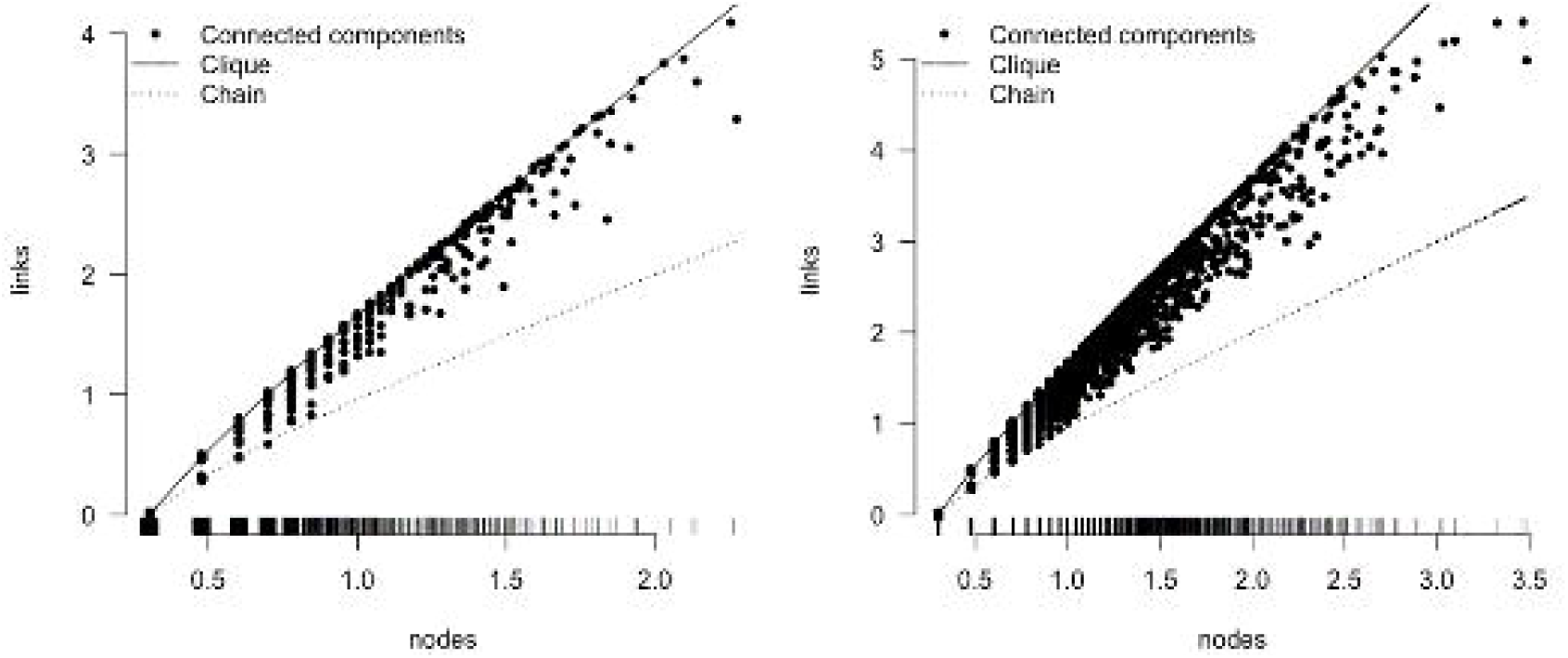
Connected components characteristics at SNP threshold equal to 12. (left: *Listeria monocytogenes*, right: *Salmonella*). Each point represents a connected component, placed on the x-axis at its number of nodes (n) and on the y-axis at its number of links, both in log 10 scale. The upper line represents n × (n - 1)/2, the maximum number of possible links and the lower, dashed line represents (n-1) the minimum number of links.

### Queries and Visualizations proposed in GenomeGraphR

GenomeGraphR provides an interactive interface where users can query, analyze, and visualize genetic connections between pathogen clinical isolates and food/environmental isolates using the curated and cleaned NCBI isolate and meta-data collection.

### Revealing connections between a set of food/environmental and clinical cases isolates

A typical use of GenomeGraphR is to reveal the existence of genetic links between a set of strains isolated from a particular source, for example from a food category, and strains from clinical cases registered in the database. [**Fig 6** presents an illustration for *S. enterica*: the set of food/environmental strains is selected from the isolation source tree by clicking, for example, on “shell eggs”. Hovering over the shell egg link, the user sees that the database includes 440 *S. enterica* strains isolated from shell eggs. The implemented query shows that, in total, 799 clinical strains are linked to 140 shell egg strains, at a SNP threshold of 12. The links between strains are structured within 52 CCs and visualized in a globe graph ([**Fig 6**). Clinical strains are represented in the form of a star and shell eggs strains as a circle, each colored by year of collection. A table listing the CCs is provided from which we can derive that connected clinical strains were collected or submitted from 1994 to July 2018, with more than 50% of strains collected or submitted after December 2016. Shell egg strains were generally collected earlier, with dates ranging from 1971 to December 2017, with a median date of January 2012. The analysis by CC shows that most strains from shell eggs were submitted before strains from clinical cases (39 out of the 52 CCs) and most of these shell eggs strains were isolated in United States (111). This suggests that it may have been possible to prevent consumer exposure if this information had been readily available. A smaller number of shell egg strains were isolated in Brazil (18), Chile (9), Australia (1) and Argentina (1). Clinical strains were isolated in the United states (691), Brazil (40), United Kingdom (39), Canada (13), Australia (9), Chile (5), Mexico (1) and Ethiopia (1). In addition to these descriptors, the user can hover over the symbol for each strain and the available metadata will be displayed. The user can also highlight the strains present in the globe graph by year, country, serovar, project center, source, or CC. The number of connected strains and their distribution within CCs depend on the chosen SNP threshold. GenomeGraphR provides users the ability to perform a sensitivity analysis, showing the changes in numbers and sizes of CCs when the threshold varies between 0 and 20 SNPs.

**Fig 6:**
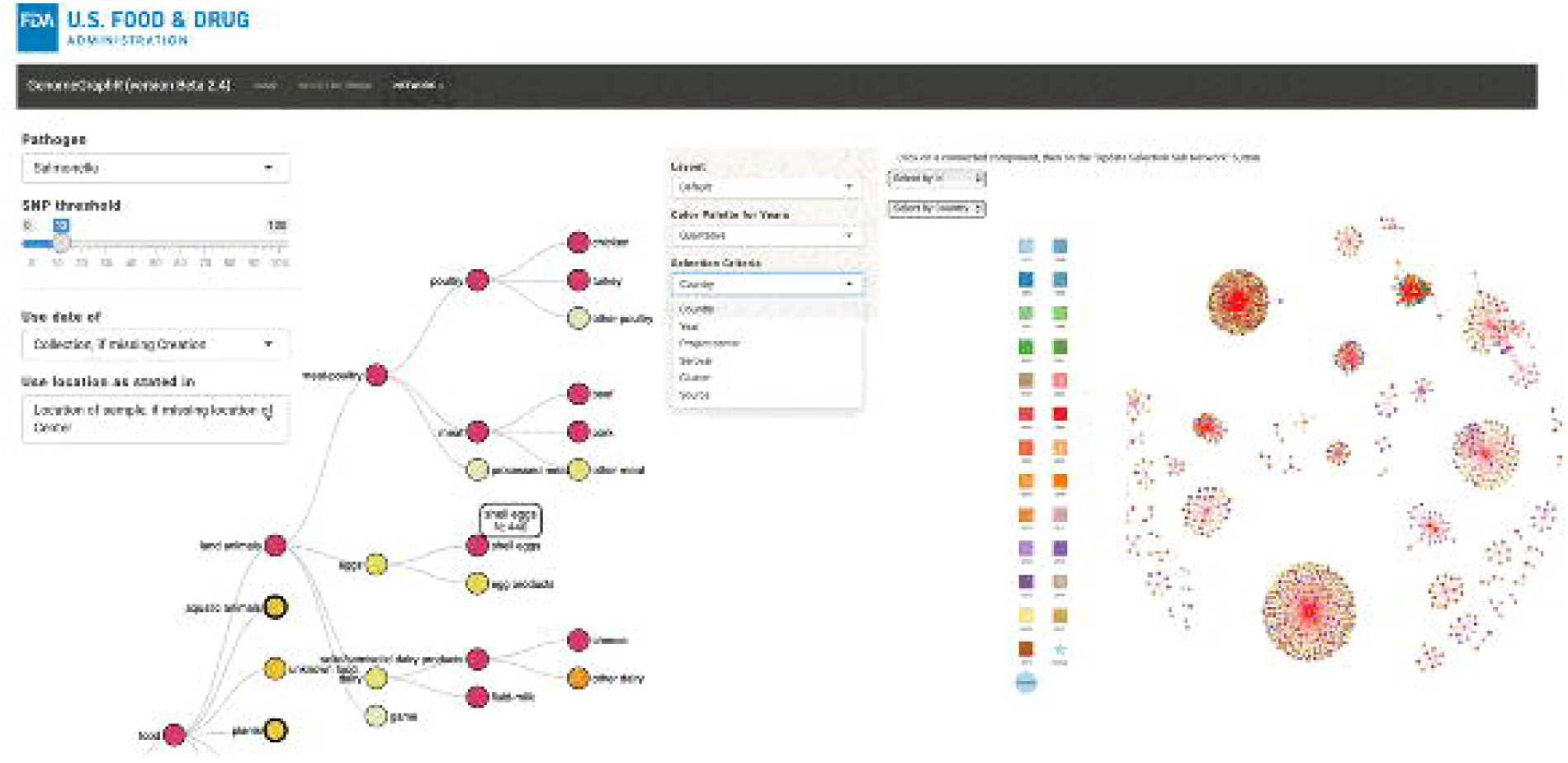
**Left: the isolation source tree.** Hovering the mouse on a node provides the number of strains from this isolation source in the database. Clicking on the node provides the graph on the right. **Right: Clinical strains connected to shell egg strains.** A connection exists when the SNP differences between a clinical strain and a non-clinical strain is less or equal to 12, leading to Connected components. The framed CC was selected to show an example of the in-depth analysis of clinical case sources.

The globe graph can be constructed for any other selection of the source category and at all hierarchical levels (Figure 7).

**Fig 7:**
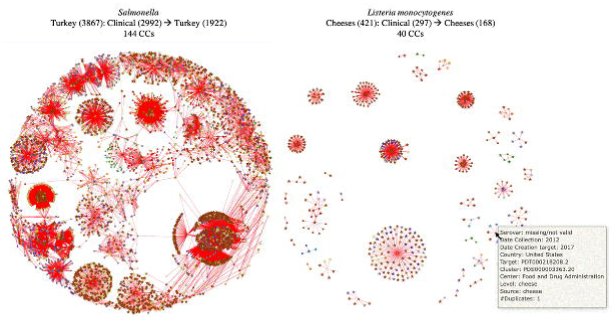
Examples of connectivity between food category and clinical cases (SNP threshold = 12). CCs: connected components, (): number of strains, ⟶ connected with SNPs ≤ 12.

GenomeGraphR provides two other ways to select a subset of strains to explore: (1) directly from the isolate table that includes associated metadata, using filtering functionalities; and (2) through a user uploaded file that contains a list of strain identifiers. From the isolate/metadata table the user can define a subset of strains by selecting criteria on one or more fields. As an example, the user may filter a subset of *S. enterica* strains isolated from Canada from food/environmental samples. This query results in 1,025 strains (Figure 8). The additional information provided within the graph show that clinical strains are primarily collected in United States.

**Fig 8:**
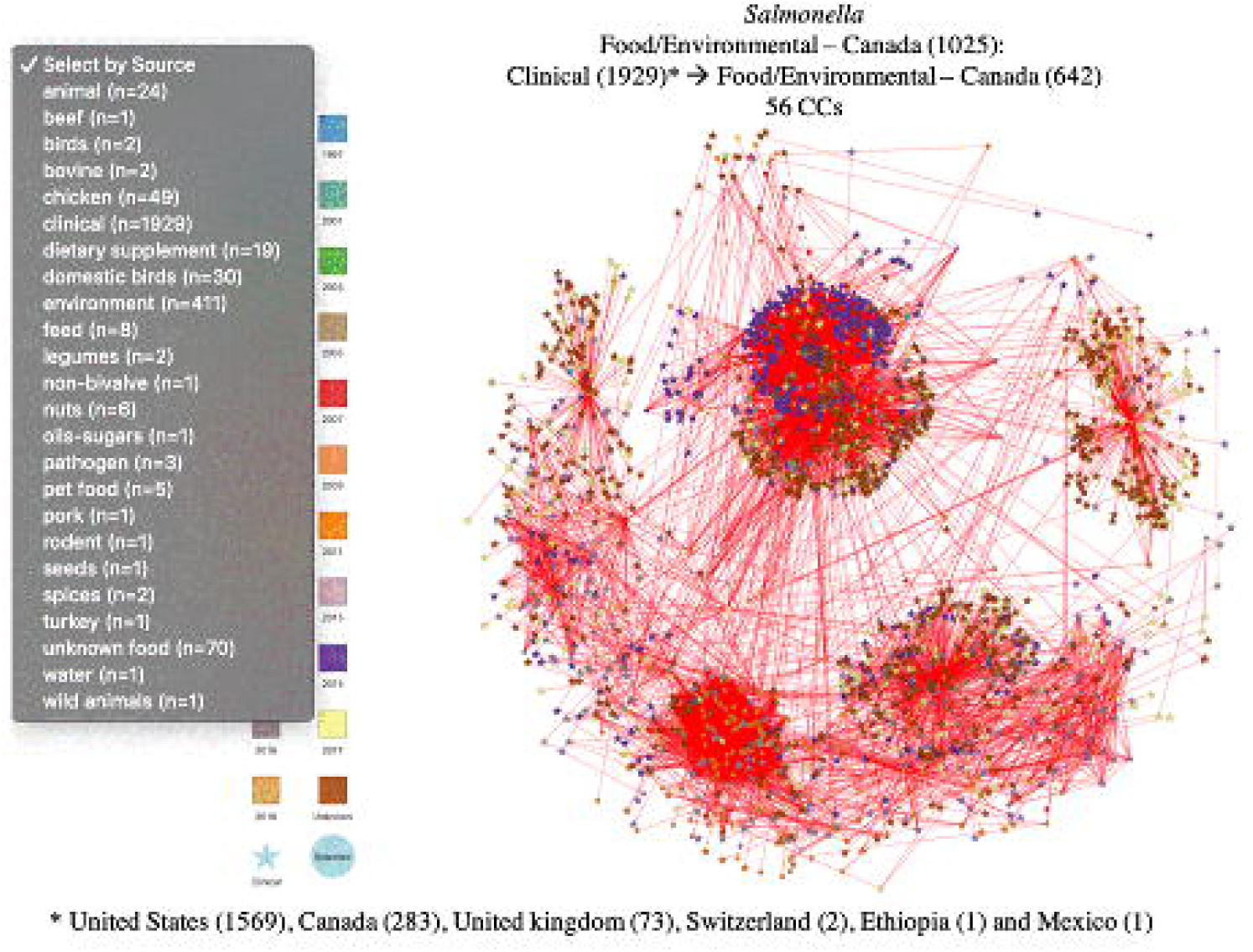
Examples of connectivity between a sub-set of strains (Food/environmental strains isolated in Canada) and clinical cases (SNP threshold = 12). CCs: connected components, (): number of strains, ⟶ connected with SNPs ≤ 12.

### Further Analysis of the revealed connections – Sub-network

As shown with the examples in [**Fig 6** - [**Fig 8**, GenomeGraphR reveals connections between clinical strains and strains isolated from food or environment (including animal and feed). These connections are grouped in CCs and each CC may represent an outbreak or more generally a group of cases associated with the same source. But the discovery of CCs linking clinical case strains with non-clinical strains (sources) is not enough evidence to attribute clinical cases to these sources. Further analysis within GenomeGraphR is needed and consists of searching in the database for the existence of other possible sources. Once the user selects a CC, GenomeGraphR runs automatically a query that searches all non-clinical strains linked to the clinical strains belonging to the selected CC. All the strains, including the CC and the new strains related to the CC clinical strains from other sources, constitute a sub-network. A specific sub-network menu provides access to features adapted to the visualization and analysis of the sub-network ([**Fig 9**).

**Fig 9:**
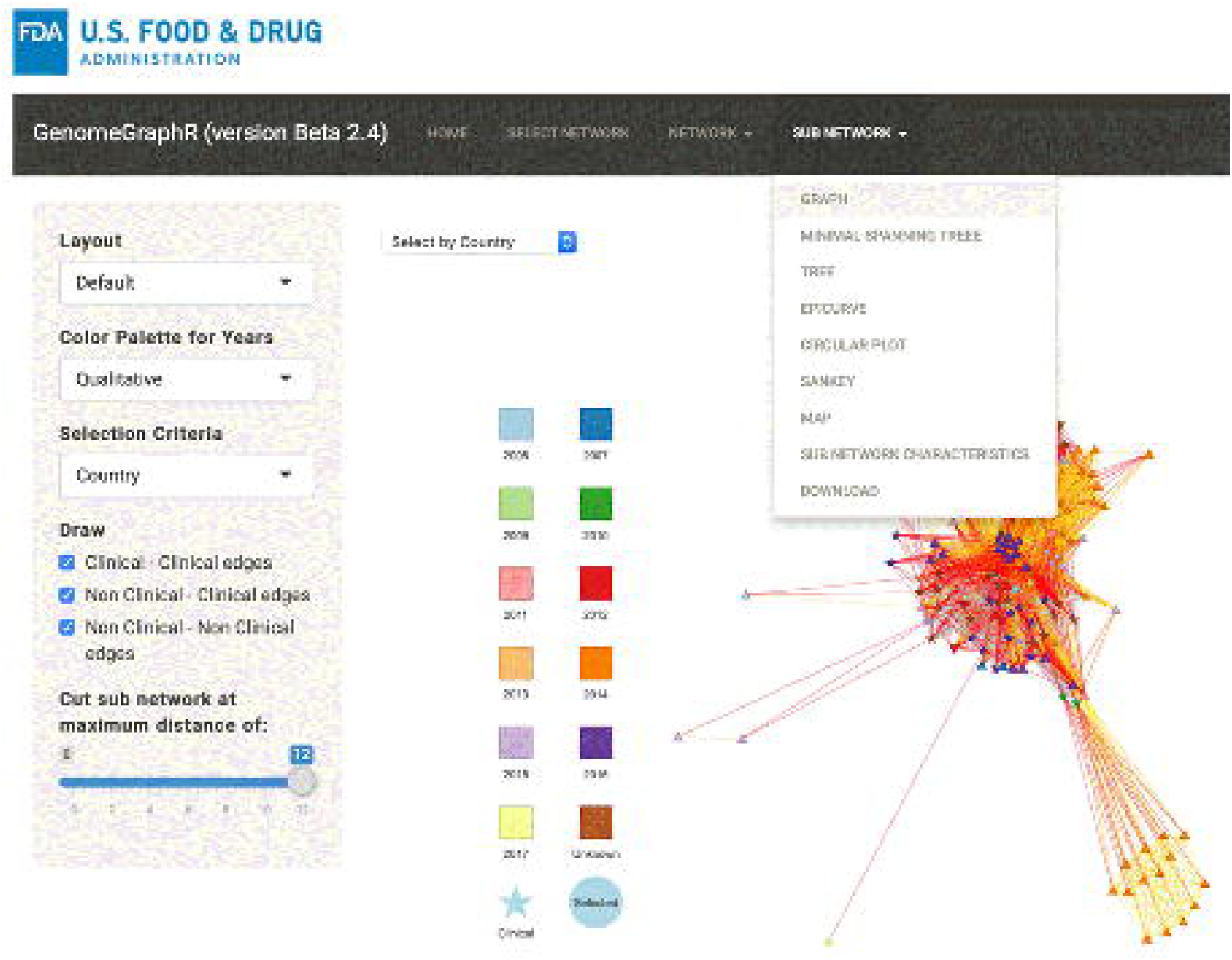
Sub-network menu.

[**Fig 10** shows an example of a sub-network associated with a CC containing 52 clinical strains and 5 strains isolated from cucumbers. (Note: this is not the CC associated with the 2015 U.S. outbreak of illness from *Salmonella* Poona in/on cucumbers; that CC is nearly symmetrical signifying little complexity and a likely single source). In addition to the CC strains, the sub-network includes 73 strains from other sources connected with the clinical strains: from water (52), environment (12), chicken (3), birds (2), animals (2), beef (1) and tomato (1). MST, circular plot and epicurve ([**Fig 10**) are valuable tools available in GenomeGraphR that help to clarify the connections between sources and clinical cases. In addition, by reducing the SNP threshold, for example from 12 to 6, or breaking the MST tree to a SNP threshold of 6, GenomeGraphR reveals three distinct clusters: two involving cucumbers and environment isolates (including water) and one involving tomato and other environment isolates including water (see supplemental materials). The dates of isolation of the strains from the two first clusters suggest that the clinical cases could be potentially linked to cucumbers contamination and indirectly to the environment from irrigation water. However, for the third cluster, the strains isolated from tomato were isolated in 2005, almost 10 years before the occurrence of the clinical cases. Surprisingly, in the cluster, a strain very close to the strain isolated from tomato (1 SNP difference) continues to be isolated in the environment and to be linked to more recent clinical strains.

**Fig 10:**
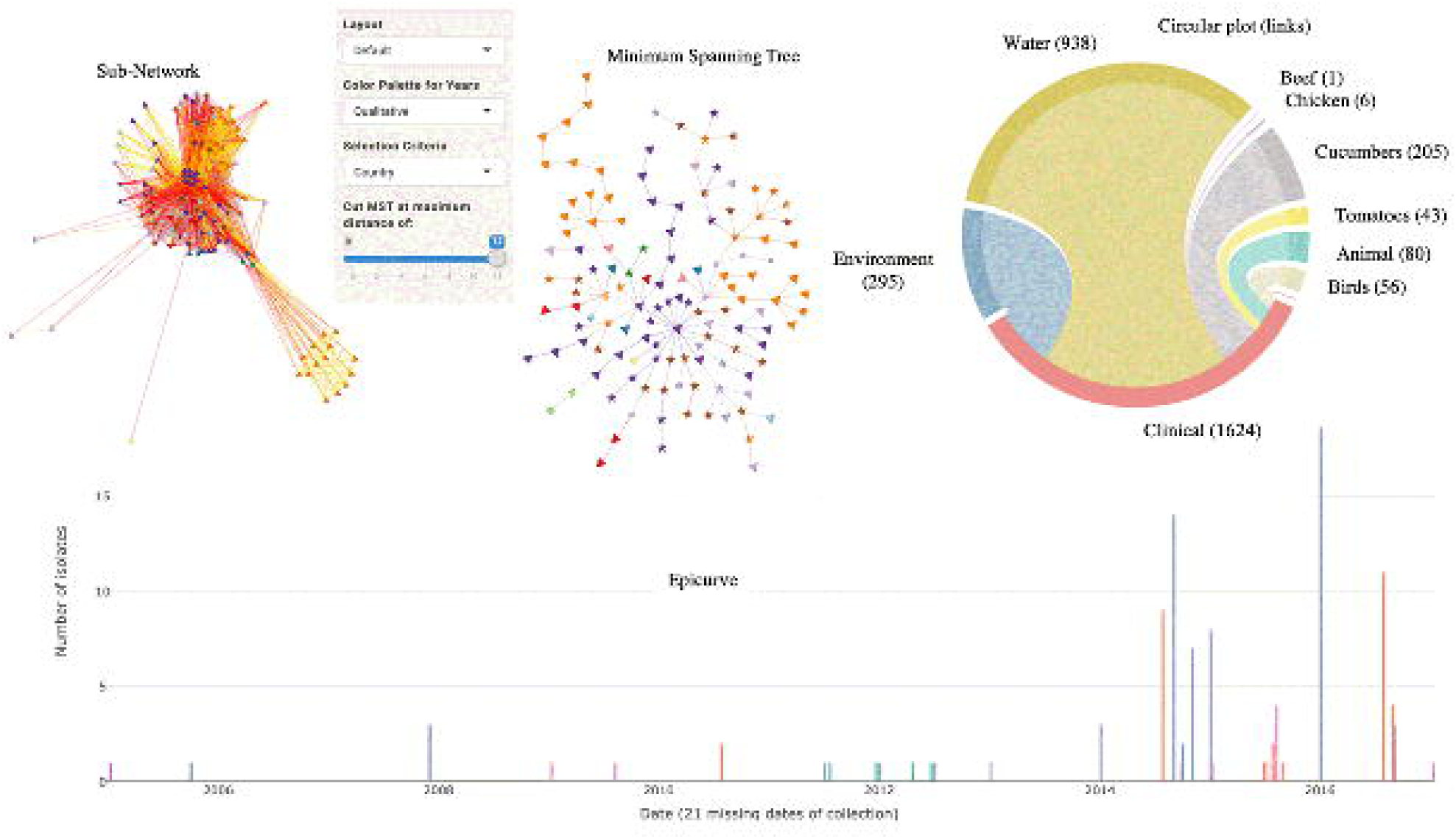
Example of Visualization and analysis of a sub-network.

[**Fig 11** provides an illustration of the map feature. From the globe graph ([**Fig 8**), which collected the links between strains collected in Canada and clinical cases in and outside of Canada, we selected one of the CCs and created a sub-network including other Food/Environment strains collected outside of Canada. The map ([**Fig 11**) shows non-clinical strains isolated from both US and Canada. Because CDC does not communicate the specific geographical locations of clinical cases, cases from the US are projected in a square area in the Atlantic. Most non-clinical strains were isolated from chicken (269) and environment (408), respectively, in Canada and the US. The map is dynamic, allowing the user to visualize the strains occurrences over time. Some of the CC strains were isolated as far back as 2001, however most strains were collected or submitted between 2014 and 2018. The MST, with SNP cut at 5, shows three main clusters: one specific to Canada, another specific to US and the last including strains from both countries ([**Fig 11**).

**Fig 11:**
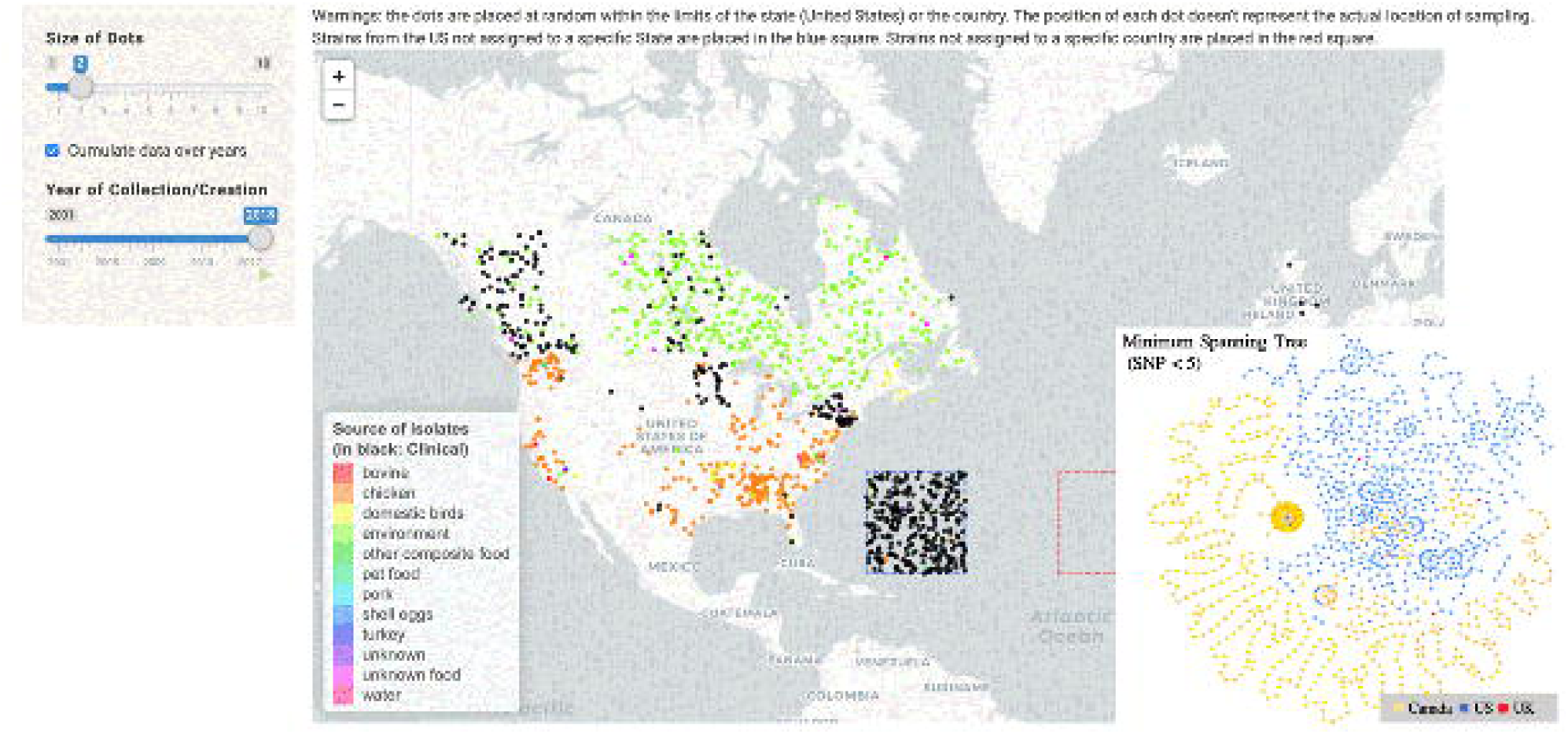
Map illustrating the origin of the strains. Note that the dots are placed at random within the limits of the state (United States) or the country (The position of each dot doesn’t represent the actual location of sampling). Strains from the US not assigned to a specific State are placed in the blue square. Clinical strains are in black.

## Discussion

To date efforts in WGS networks have focused primarily on how to discover the most complete and accurate set of nucleic acid sequences for outbreak identification [1]. This scope of outbreak identification can be fulfilled with non-standard source metadata, as a relatively small set of strains are considered for a given outbreak. But the integration of WGS data with high quality epidemiological and food chain data is crucial for other uses of these databases, notably for larger scale analyses [20].

In our study, we developed and applied procedures to cleanup and further organize the NCBI Pathogen Detection metadata to facilitate its use for population level epidemiology studies and risk assessment. With WGS data growth, metadata verification and validation procedures will need to be applied frequently to ensure GenomeGraphR data are comprehensive and up-to-date. The processes of cleanup, organization, and integration can be regularly applied, as we have done here; reducing missing and ambiguous data will require commitment of NCBI submitters to complete accurately all the mandatory fields and the NCBI Pathogen Detection pipeline to provide enhanced standardized options and controlled vocabularies for data input. Addition of alternative isolation source classification schemes in GenomeGraphR is anticipated, *e.g*., schemes being developed by the GenEpiO consortium (http://genepio.org/home/consortium/) and schemes used in other countries or regions, such as FoodEx2 (European Food Safety Agency (EFSA) food classification and description system) and will be facilitated by improvements in metadata quality.

Various applications exploiting WGS [22] have been developed, but we believe that none of them fully meets our needs for research and analysis in the context of food safety. The purposes of the currently available applications, such as Microreact (https://microreact.org), Nextstrain (https://nextstrain.org) or tools available within the Integrated Rapid Infectious Disease Analysis (IRIDA) project platform (www.irida.ca) are mainly to aid epidemiological understanding of non-foodborne pathogen spread and evolution, especially in outbreak situations. Moreover, these tools primarily use phylogenetic trees to visualize WGS data. As datasets become larger and trees increase in size, visualizing a tree with thousands of branches or leaves becomes a scalability issue, and other solutions are needed to provide alternative visualizations [23]. PHYLOViZ Online [24], one of the available tools in IRIDA platform, offers a scalable methodological alternative to traditional phylogenetic inference approaches: MST analysis of allelic data used in epidemiological investigations and population studies of bacterial pathogens. More recently, GrapeTree was made available [25]. It was designed to facilitate reconstruction and visualization of genomic relationships of large numbers of bacterial genomes based on a novel minimum spanning tree algorithm (MSTree V2). GrapeTree is integrated into EnteroBase (http://enterobase.warwick.ac.uk/) and provides an alternative to the classical phylogenetic trees when the number of genomes is large with interactive integration of the available metadata.

These existing applications and tools provide solutions to overcome significant computational challenges in terms of genomics result analysis and reporting. We have built on these innovations in GenomeGraphR, where appropriate and possible, and added new analytic strategies, including network analysis visualizations and flexible and robust search features.

When using GenomeGraphR for population level epidemiology studies or risk assessment, we are faced with the issue of dataset collection bias; the collection of strains available in the NCBI database is not *a priori* representative of all strains circulating in the food chain. While we have demonstrated that the probability for a new strain to be “close” to a strain already present in the database (even within a 50 SNP threshold) is increasing with time (presumably associated with the increase in volume and quality of WGS data), this probability is still low. Application and integration of appropriate statistical methods to address collection bias are needed to ensure accurate interpretation of the relationships revealed in GenomeGraphR.

Despite these challenges, by combining data cleaning and enhanced organization with analytic and visualization tools, we were able to reveal connections between foods or food categories and cases and describe the distribution of these in time and space.

## Conclusion

GenomeGraphR is a unique flexible application, that presents foodborne pathogen WGS data and their associated metadata designed to address a variety of research questions relating to, for example, transmission sources and dynamics, global reach, and persistence of genotypes associated with contamination in the food supply and foodborne illness across time or space. With the development of specific analytics to address missing values, bias, and non-representative sample/strain, these data may provide novel approaches to foodborne illness attribution at the population level and will provide critical data for exposure assessment and hazard characterization (dose-response) needed for risk assessment. Integration of isolate genome characteristics in GenomeGraphR in the future, including anti-microbial resistance, virulence, and persistence factors, will allow further discernment among genetically related strains, and will enhance the scope of research insights available from WGS integration for population level epidemiology research and food safety risk assessment.

## Acknowledgments

We thank Marc Allard and Ruth Timme for their review and comments on a draft of this manuscript. We thank Sherri Dennis for her support of this work throughout its development. This work fulfills in part recommendations from the Interagency Risk Assessment Consortium Workgroup, “Exploration of the Implications of Whole Genome Sequencing on the Conduct and Application of Risk Assessment in Food Safety Decision-Making.”.

